# Silencing of a unique integrated domain nucleotide-binding leucine-rich repeat gene in wheat abolishes *Diuraphis noxia* resistance

**DOI:** 10.1101/261644

**Authors:** Vittorio Nicolis, Eduard Venter

## Abstract

Plants respond in a similar manner to aphid feeding as to pathogen attack. *Diuraphis noxia* is a specialist aphid, feeding only on selected grasses that include wheat, barley, and oats. The wheat-*Diuraphis noxia* interaction is characterized by very similar responses as seen in wheat-pathogen interactions with none of the underlying resistance pathways and genes characterized yet. From wheat harboring the *Dn1* resistance gene, we have identified a nucleotide-binding leucine-rich repeat (NLR) gene containing two integrated domains (IDs). These are three C-terminus ankyrin repeat-domains and an N-terminus WRKY domain. The NLR core of the gene can be traced through speciation events within the grass family, with a recent WRKY domain integration that is *Triticum* specific. Virus induced gene silencing of the gene in resistant wheat lines resulted in the abolishment of localized cell death. Silenced plants supported a higher number of aphids similar to the susceptible NIL and the intrinsic rate of increase of the aphids matched that of aphids feeding on the susceptible NIL. The presence of the gene is necessary for *Dn1* resistance and we have named the gene *Associated with Dn resistance 1* (*Adr1*) to reflect this function.

## Introduction

The Russian wheat aphid *(Diuraphis noxia* Kurdjumov) is a specialist aphid pest of grasses. Its primary hosts with commercial importance are wheat, barley, and oats, while it can survive well on false barley, wild oats, and rescue grass (Jankielsohn 2013). After introduction and escalation into a pest, *D. noxia* causes tremendous losses in wheat production countries (Morrison and Peairs 1998). Resistant wheat cultivar development resulted in lower yield losses being incurred, but also increased pressure on the aphids to develop new biotypes. To date twelve resistance genes (*Dn*-genes) have been identified and employed in developing resistant cultivars. Several of these genes were incorporated into resistant wheat lines in South Africa and this has led to the development of at least four known aphid biotypes (RWASA1 – 4) that overcame all but the *Dn7* resistance gene (Jankielsohn 2016). The *Dn1* resistance gene, effective against RWASA1, has an antibiotic effect against the aphids and limits their fecundity, growth and longevity (Smith et al. 1992). This gene was the first gene to be implemented in resistant wheat breeding and maps to chromosome 7DS (Bierman 2015). The gene has yet to be identified or cloned and the mechanism by which it contributes resistance is not clear.

Plants under attack by phloem feeding insects respond similarly to attack by pathogens (Bos et al. 2010; Rodriguez and Bos 2013). *Diuraphis noxia* feeding progresses through probing on the leaf surface, intercellular stylet navigation and finally, penetration into the phloem to feed. During the intercellular phase the stylet occasionally pierces cells for the aphid to sense its way. During transition through the leaf, the stylet is protected by the production of a salivary sheath that encloses the stylet. This sheath consists of saliva that solidifies once it is secreted from the stylet tip. Once the stylet reaches the phloem the aphid starts to produce watery saliva that assists in feeding and that contains effector molecules that interfere with the plant’s defense responses (Bos et al. 2010; Lapitan et al. 2007; Rodriguez and Bos 2013; Will et al. 2007). Indeed, it has been shown by Lapitan et al. (2007) that the injection of proteins from different salivary fractions induces symptoms in a susceptible cultivar. This included chlorosis visualized as chlorotic lesions and streaking, leaf rolling, and stunted growth. In contrast, the resistance response is likened to the classic gene-for-gene interaction mediated by an R-gene with hypersensitive-linked necrotic lesions observed at the site of aphid feeding (Botha et al. 2005).

Nucleotide-binding leucine-rich repeat (NLR) genes often work as dimers to form receptor complexes to specify resistance against a pathogen (Sinapidou et al. 2004). In cereals, several NLR-complex receptors have been identified, with Lr10 and RGA2 the first identified from wheat (Loutre et al. 2009). Studies on rice blast resistance alleles indicated that resistance is imparted by a combination of two genes located at the same locus, e.g. the *Pikm* and *Pia* loci (Ashikawa et al. 2008; Okuyama et al. 2011). Thus, interaction between NLRs are indicative of a complex avirulence effector-recognition system that plants employ during innate immunity. Interaction between NLRs to provide resistance can include up to three proteins working in conjunction as seen in barley resistance against *P. graminis.* Here, the NLRs *rpg4* functions together with *Rpg5, HvRga1,* and *HvAdf3,* an actin-depolymerizing factor-like gene (Wang et al. 2013). RPG5 contains an additional serine/threonine protein kinase domain that could implicate this NLR in signal transduction. Thus, pathogen recognition by the RPG5/HvRGA1 complex may initialize signal transduction by the phosphorylation of serine/threonine protein kinase domains (Wang et al. 2013). This was postulated after the observation that the RPG1 resistance gene in barley contains a similar serine/threonine protein kinase domain that is rapidly phosphorylated in the resistant line, against *P. graminis* f. sp.*tritici,* but not in the susceptible line (Nirmala et al. 2010). This domain could also be targeted by the rust effector and is guarded by the NLRs that are bound to it. Decoy domain recognition has been demonstrated by et al. (2015) and Le Roux et al. (2015) where the RPS4/RRS1 dimer interacts directly with effectors, PopP2 and AvrRps4, via an integrated WRKY domain to induce a defense response.

Nucleotide-binding leucine-rich repeat genes that contain WRKY integrated domains (NLR-ID) at their C-terminus have been identified to act as decoy binding sites for interaction with effector proteins (Le Roux et al. 2015; Sarris et al 2015). RRS1-R and RPS4 impart resistance to *Arabidopsis* against *Ralstonia solanacearum* and *Pseudomonas syringae* pv. *pisi* (Sarris et al. 2015). PopP2 is an acetyltransferase that specifically acetylates the lysine residues located in the WRKYQK motif of RRS1 and other nuclear localized WRKY transcription factors that then interferes with their DNA binding capability. Thus, disabling transcription activation and subsequent defense responses. Acetylation disrupts the DNA binding of RRS1-R and in turn activates RPS4-dependent resistance by releasing the RRS1/RPS4 complex inducing innate immunity in *Arabidopsis.* Thus, turning the pathogen’s effector against itself to induce innate immunity in the cell (Le Roux et al. 2015).

The identification of NLR-ID decoy proteins that dimerize to target effector proteins has furthered our understanding of the complexity of plant innate immunity against pathogens. Identification of a similar NLR-ID protein from wheat that is implicated in the defense response to *D. noxia,* has led us to functionally test the role that it plays in the wheat-*D. noxia* interaction. We hypothesized that *5AL-B4,* a C-terminal WRKY containing NLR-ID with additional N-terminal ankyrin repeats, plays a role during the innate immunity of wheat carrying the *Dn1* resistance gene against *D. noxia*. It is intriguing that *5AL-B4* contains integrated WRKY and ankyrin domains, as ankyrin, WRKY10, −12, and −53 transcription factors have previously been associated with *D. noxia* resistance (Smith et al. 2010; Van Eck et al. 2010). Therefore, the aims were to further characterize *5AL-B4* and to use VIGS-mediated silencing to ascertain the role that *5AL-B4* plays during wheat interacting with *D. noxia.*

## Results

### *5AL-B4* homologs are genetically similar

TRIAE_CS42_5AL_TGACv1_374266_AA1195550 (November 2014 *T. aestivum* Ensemble release; designated *5AL-B4)* is an NLR with similar architecture to NLR-ID decoys against pathogen effectors (Fig. 1). It shared homology with *Pi36* and identified as a possible role player in the wheat-*D. noxia* interaction from the study by Nicolis et al. (2017). The sequence and domain architecture for *5AL-B4* was predicted from cv. ‘Chinese Spring’, which has no resistance to *D. noxia.* PCR amplification and sequencing of *5AL-B4* from a *D. noxia* susceptible near isogenic line (NIL) Tugela, and the resistant NIL Tugela DN, was used to confirm the presence of predicted domains and to search for SNPs between the resistant and susceptible NILs. No SNPs were identified between the resistant, susceptible, and Chinese Spring cultivars and all predicted domains from Chinese spring were preserved throughout both NILs. These predicted and confirmed domains included three N-terminal ankyrin repeats followed by a CC-NB-LRR architecture with a single C-terminal WRKY domain (Fig. 1A). The position of the WRKY domain in 5AL-B4 is consistent with the recently postulated NLR-ID genes. This position is further consistent with the recently characterized function of RRS1 that interacts with PopP2 and AvrRps4 (Le Roux et al. 2015; Sarris et al. 2015).

**Fig. 1.**
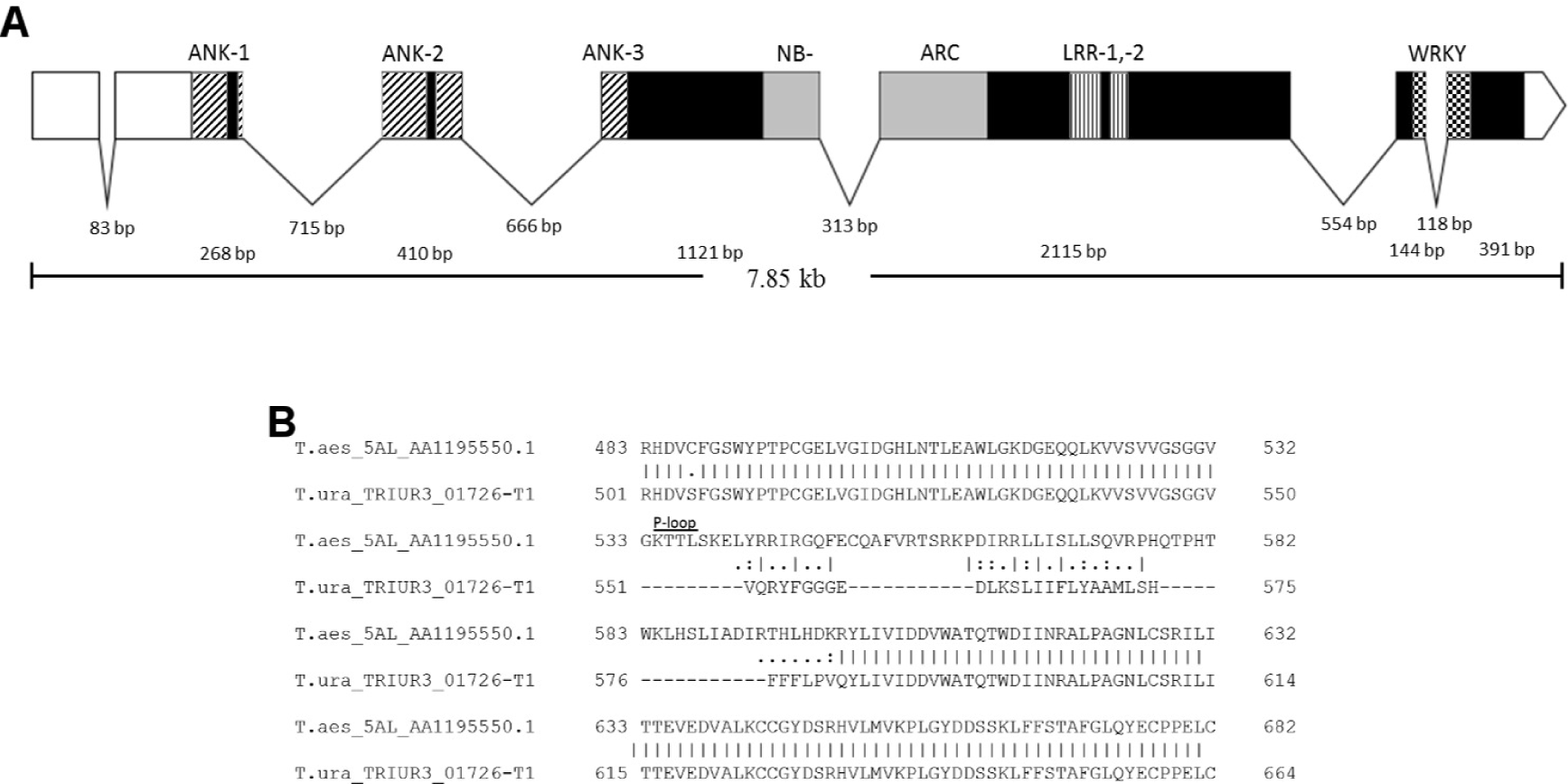
Structure of *5AL-B4.* **A,** *5AL-B4* encodes seven exons and six introns with the predicted untranslated regions (empty boxes), ankyrin, NB-ARC, LRR, and WRKY domains indicated. **B,** Sequence alignment between 5AL-B4 and ancestral copy in *T. urartu* indicating sequence dissimilarity around the P-loop.

### *5AL-B4* is regulated during the wheat-*D. noxia* interaction

The possible differential regulation of *5AL-B4* in RWA-SA1 infested resistant NIL Tugela DN and susceptible NIL Tugela was tested using ddPCR. The use of droplet digital PCR (ddPCR) for transcript level determination was due to the low levels of NLR expression reported rendering amplification to establish a standard curve for RT-qPCR analysis not consistent. The low levels of NLR expression is known and hinders expression analysis (du Preez et al. 2008; Fossdal et al. 2012). Variation was detected in both NILs for the expression of *5AL-B4* from nine time points (Fig. 2). In the susceptible Tugela, only significant (p = 0.023) downregulation was detected from the early to late time points. Downregulation occurred for 0-4 hpi (p =0.031), 0-48 hpi (p = 0.023), 1-48 hpi (p = 0.05), 4-48 hpi (p = 0.05), and 6-48 hpi (p = 0.049). In the resistant Tugela DN, significant upregulation was detected at the early time points for 0-6 hpi (p = 0.044), 0.5-1 hpi (p = 0.036), 0.5-6 hpi (p = 0.029), and 0.5-8 hpi (p = 0.036). This was followed by significant downregulation of transcript numbers for 0-24 hpi (p = 0.035), 1-24 hpi (p = 0.035), 6-24 hpi (p = 0.008), 8-24 hpi (p = 0.033), and 1-48 hpi (p = 0.05).

**Fig. 2.**
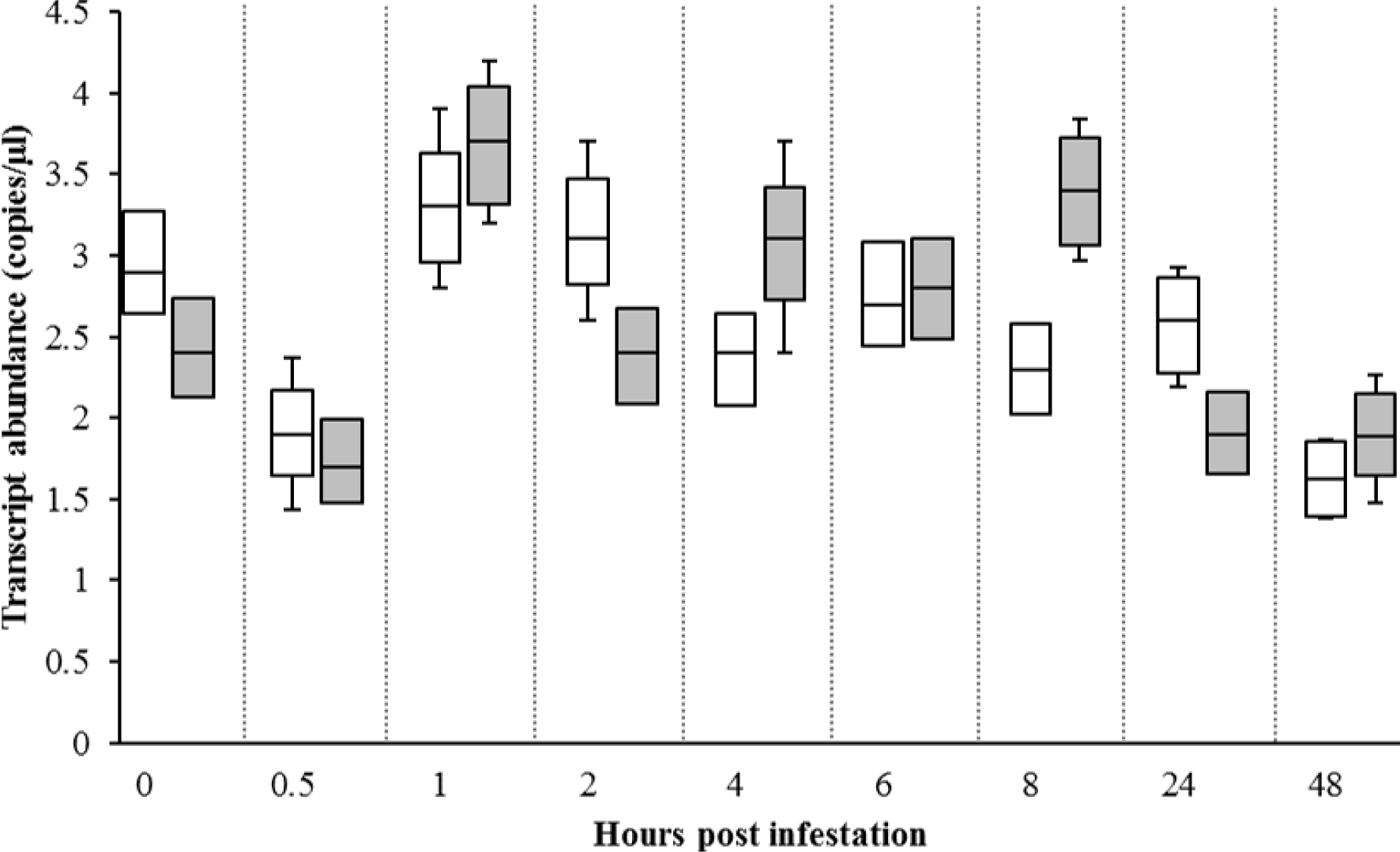
Differential expression of *5AL-B4* transcripts between Tugela (white bars) and Tugela DN (grey bars) NILs. Significant downregulation of transcript levels in the susceptible NIL Tugela was detected from the early to late time points (4-48 hpi, p ≤ 0.05). In the resistant NIL Tugela DN, significant upregulation from the early to intermediary (0-8 hpi, p ≤ 0.05) time points was detected, followed by significant downregulation at 24 and 48 hpi (p ≤ 0.05).

### A *T. urartu* NLR-ID contributed the core of *5AL-B4*

Phylogenetic analysis of the full-length protein sequences closely related to *5AL-B4* revealed that *T. urartu* TRIUR3_01726 is the closest relative to *5AL-B4* (Fig. 3). *Triticum urartu* is the donor of the A-genome in hexaploid wheat. The core sequence of TRIUR3_01726 served as a scaffold for the integration of the WRKY domain now present in *5AL-B4* (solid black arrow). The protogene of *5AL-B4* follows speciation of the pooid, oryzoid and panicoid lineages, across 40 million years, with possible duplication and divergence occurring at each speciation event. A duplication and divergence event is present where the oryzoid species diverge from panicoid and pooid species as identified by related genes in *Oryza* species (open grey arrow). Soon after, the gene in the panicoid and pooid lineages duplicated, diverged and entered the pooid clade, with *Brachypodium distachyon* occurring basal to the divergence between *Hordeum* and *Triticum* species (open grey arrow). The NLR core continued to duplicate and diversify within barley and wheat species, with barley occurring basal to each branch, including the branch containing the *5AL-B4* gene and its homoeologs TRIAE_CS42_5B-AA13L_TGACv1_405227_AA1322580 *(5BL-AA13)* and TRIAE_CS42_5D-AA14L_TGACv1_434466_AA1436530 *(5DL-AA14)* (solid grey arrows). Respectively, *5BL-AA13* and *5DL-AA14* have fused with an Apetala 2 (AP2) and a WRKY-WSKY domain independently to the *5AL-B4* WRKY fusion. The function of the NLR core in defense response would seem to have been crucial and therefore maintained within this clade. However, there must be an advantageous selective pressure that exists on the core NLR recruiting additional domains, i.e WRKY, WRKY-WSKY, and the AP2 domains. The recruiting of multiple domains to the core NLR happened independently in the three homoeologous genomes of *T. aestivum* possibly through convergent evolution to increase the relevant function of each homolog. As these domains all represent transcription factors it is tempting to speculate that these homoeologs might be guarding transcription factors that are targeted by effectors from pathogens and pests.

**Fig. 3.**
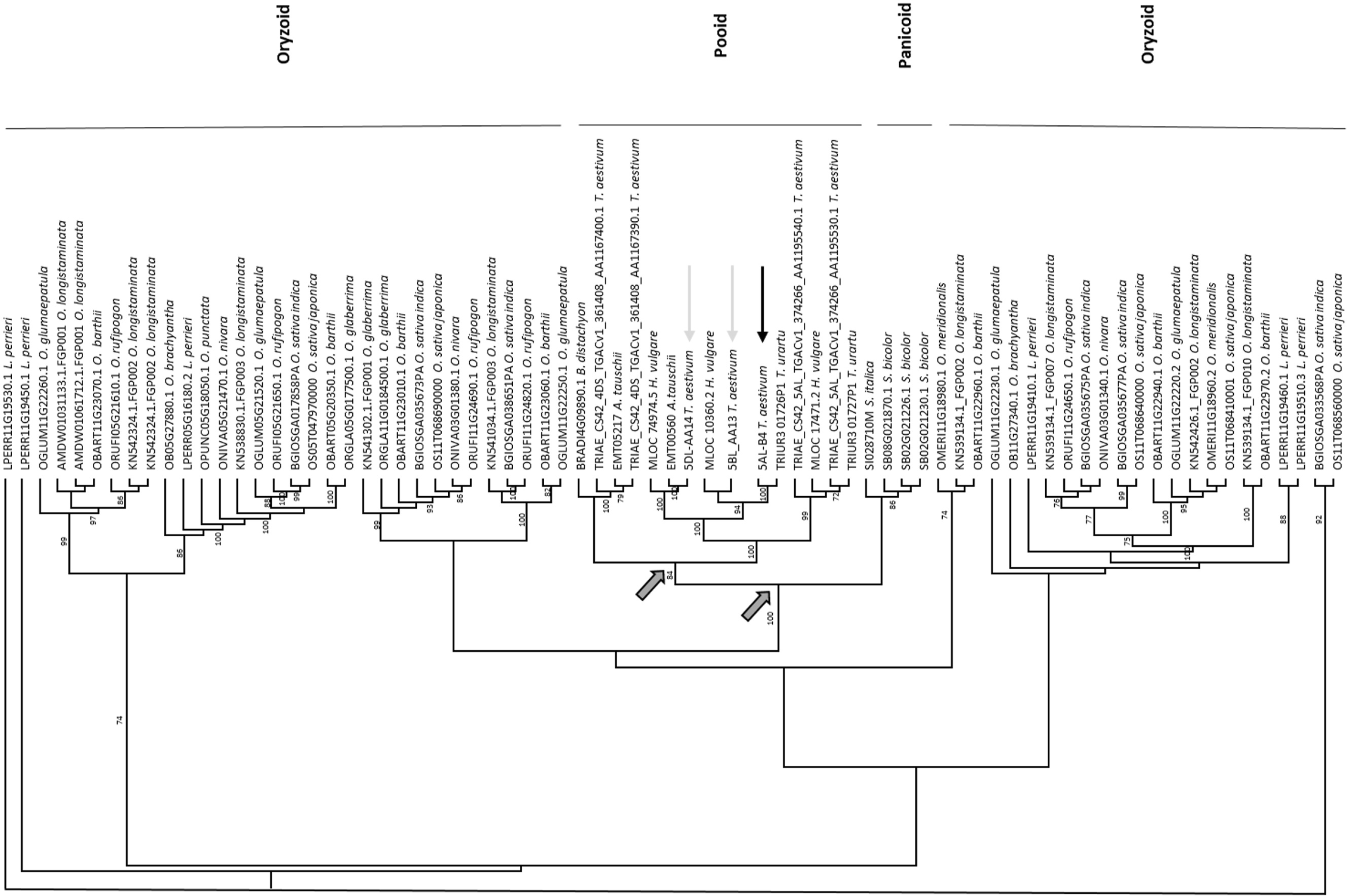
Phylogenetic analysis indicated that the ANK-domain containing core sequence of *T. urartu* TRIUR3_01726 served as a scaffold for the integration of the WRKY domain of 5AL-B4. The most parsimonious tree is presented with a consistency index of 0.7189 and retention index of 0.7841. Bootstrap values (1,000 repetitions) above 70 *%* are indicated. Solid black arrow indicates *5AL-B4,* solid grey arrows indicate the two homoeologs *5BL-AA13* and *5DL-AA14.* The split between the *Triticum, Hordeum* and *Oryzae* clades are indicated with open grey arrows.

### TaWRKY50, the most likely candidate for *5AL-B4* integration

To further understand the integration of a WRKY domain onto the NLR *5AL-B4*, we investigated the phylogenetic placing of the *5AL-B4* specific WRKY domain. All publically available *T. aestivum* WRKY domain amino acid sequences were aligned to the WRKY and WRKY-WSKY domains on the two homoeologs occurring on the 5AL and 5DL chromosomes (Fig. 4). This analysis indicated that the integrated WRKY domain on the *5DL-AA14* (solid grey arrow) homoeolog is a perfect match to TaWRKY41. This integration event conforms to a group I WRKY domain, with two WRKY domains occurring in close proximity of each other. Thus, the core NLR recruited its WRKY domain from the active transcription factor TaWRKY41. Surprisingly, there is no matching TaWRKY domain that corresponds to the integration in *5AL-B4* (solid black arrow), with the closest match being TaWRKY50 at 79.7 *%* shared homology across the studied 69 amino acids comprising the domain. The WRKY that was integrated onto *5AL-B4* also originated from group III WRKYs that are characterized by additional amino acid inclusions and the occurrence of C2HC zinc finger conformation at the N-terminal end (Eulgem et al. 2000). Interestingly, the two domains on 5DL-AA14 contain the two group I WRKY with the first being a WRKY motif. However, it also contains the extra amino acids that are present in the group III WRKYs that results in the grouping of the second WSKY motif with group III WRKYs.

**Fig. 4.**
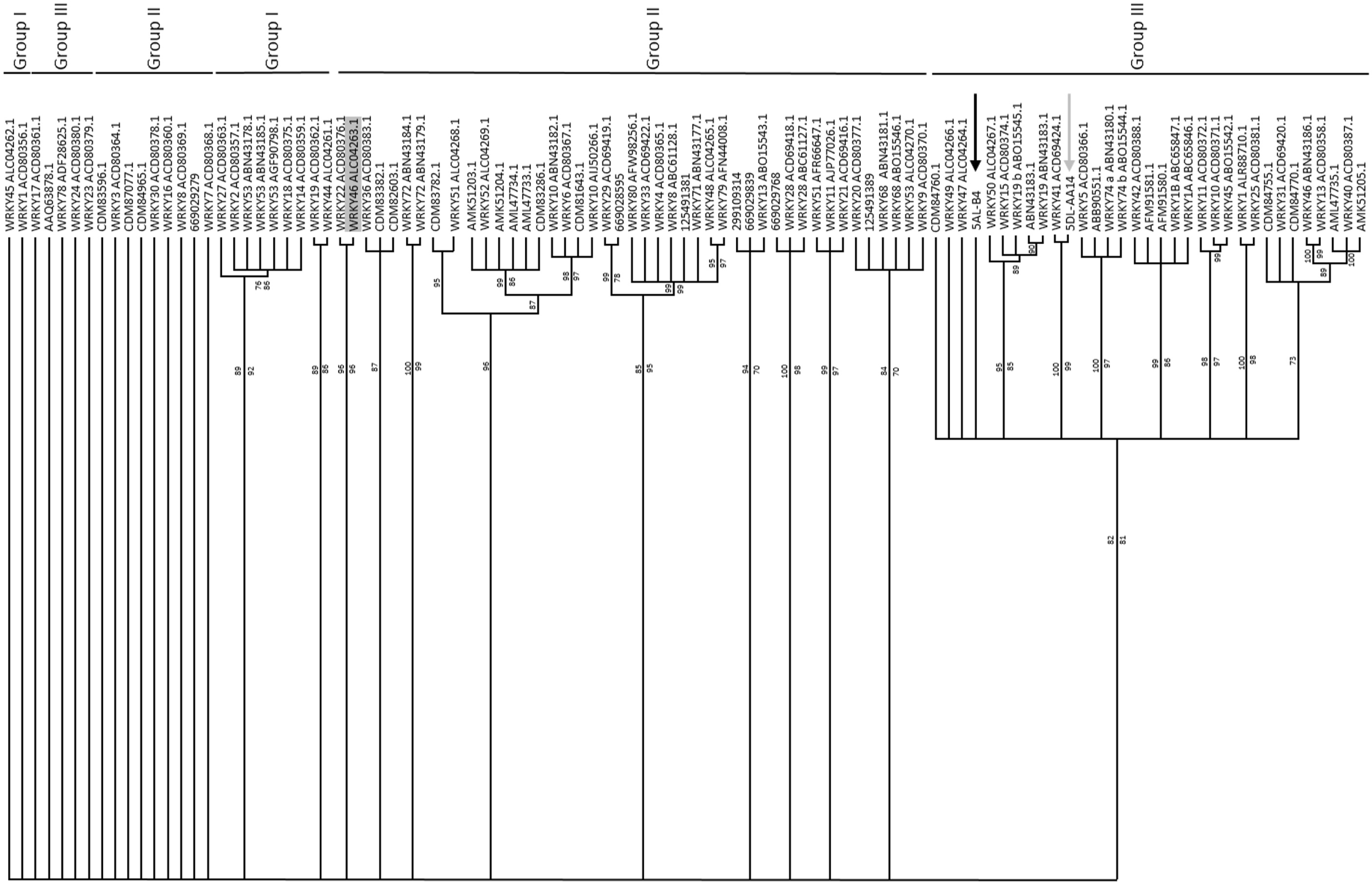
*Triticum aestivum* WRKY-domain containing sequences. The most likely candidate WRKY domain that was integrated into 5AL-B4 is TaWRKY50. Multiple WRKY-domain integrations, spanning all the sub-groups, are evident in contrast to the proposed ancient integration of AtWRKY46. The most parsimonious tree is presented with a consistency index of 0.5530 and retention index of 0.8530. Bootstrap values (1,000 repetitions, > 70 %) above branches for parsimony analysis and below for maximum likelihood analysis. Best maximum likelihood model fit DCMut +F. Solid black arrow indicates *5AL-B4,* solid grey arrow indicate the WRKY-containing homoeolog *5DL-AA14.*

### Multiple integration events since monocotyledonous divergence

It was postulated that the integrated WRKY fusions in monocotyledonous species occurred prior to the divergence between panicoid, pooid, and oryzoid species and involved a WRKY46 homolog (Sarris et al. 2016). Analysis of the integrated WRKY domains from a representative number of monocotyledonous NLR-IDs reveals that integration of the WRKY domain is host and lineage specific (Supplementary Fig. S1). Indeed, the various integration events were not based on WRKY46 (highlighted) and involves unrelated WRKY homologs. For the majority of the integration events the NLR-ID WRKY did not diverge extensively from the WRKY that it could have originated from. However, in the case of *5AL-B4* this is not evident (Fig. 4). Analysis indicated that the closest WRKY to the one found on *5AL-B4* is TaWRKY50, but with extensive divergence from the original sequence.

### The effect of *5AL-B4* transcript silencing on aphid and plant performance

The role of *5AL-B4* in the wheat-*D. noxia* interaction was further evaluated using barley stripe mosaic virus (BSMV)-based gene silencing (VIGS). The expression of *5AL-B4* was knocked down in resistant Tugela DN using a unique gene specific sequence following the WRKY domain. Absolute quantification using ddPCR was used to determine the levels of *5AL-B4* silencing achieved by VIGS compared to transcript levels in the susceptible and resistant controls. BSMV treatment reduced the expression levels of *5AL-B4* by 40 *%* compared to the uninoculated susceptible and resistant controls (Fig. 5C). Levels of *5AL-B4* were slightly elevated in the TDN+BSMV0 treatment, mostly likely due to the stress of viral infection. These levels of silencing were similar across three replications of the VIGS experiments and similar to those observed in other VIGS studies for non-NLR targets (Schultz et al. 2015; Senthil-Kumar and Mysore 2011).

The reproduction of aphid foundresses in individual clip cages was monitored in order to determine the effect of silencing *5AL-B4* on *D. noxia* RWASA-1 performance (Table 1, Fig. 5D). The average number of nymphs born per day on TDN+BSMV5AL-B4 (2.47 nymphs day^-1^) were similar to the 2.70 nymphs day^-1^ on the susceptible Tugela control and were significantly (p = 0.012) more than in the resistant controls Tugela DN (1.71 nymphs day^-1^) and TDN+BSMV0 (1.79 nymphs day^-1^). The average total number of nymphs produced can be used as a measure of aphid fertility (Van Eck et al. 2010). This showed that after 16 days of feeding (21 days post viral infection), a mean total of 22.3 offspring had been produced on TDN+BSMV5AL-B4 plants with a similar number of offspring (24.3) observed on Tugela plants. In contrast, the resistant controls Tugela DN and TDN+BSMV0, produced 15.4 and 16.1 mean total offspring respectively. The intrinsic rate of increase (*r*_m_) was used as a measure of aphid fecundity for the different treatments. The highest calculated rate of increase was observed in aphids feeding on susceptible Tugela plants (0.302) and aphids feeding on the resistant control Tugela DN had the lowest rate of increase (0.252). This was not significantly different to the rate of increase on the TDN+BSMV_0_ control (0.258). Aphids feeding on TDN+BSMV5AL-B4 had a significantly higher rate of increase (0.295) compared to the Tugela DN and TDN+BSMVo controls (p = 0.000011). The prenymphipositional period was recorded for all treatments, however there was no significant difference in the number of days from the birth of the foundress to the start of her reproduction between the four treatments. The start of reproduction was on average seven days after the birth of the foundress, which is consistent with other VIGS studies performed on this interaction (Anderson et al. 2014).

**Table 1.** Aphid fecundity on silenced and control plants. Plants treated with BSMV_*5AL-B4*_ had a similar mean aphid production rate and aphid fecundity to the susceptible control, and significantly more than the resistant controls. No difference was observed in the prenymphipositional period for each treatment. Different letters per column indicate statistical significance between values listed in that column.

| Treatment | Mean births day<sup>-1</sup> | $r_m$ | Prenymphipositional period (days) |
| --- | --- | --- | --- |
| Tugela | 2.70a | 0.303+0.021a | 7.5+0.5a |
| TDN+BSMV<sub>0</sub> | 1.79b | 0.258+0.013b | 7.6+0.5a |
| Tugela DN | 1.71b | 0.252+0.012b | 7.8+0.4a |
| TDN+BSMV<sub>5AL-B4</sub> | 2.47a | 0.295+0.02a | 7.4+0.5a |

**Fig. 5.**
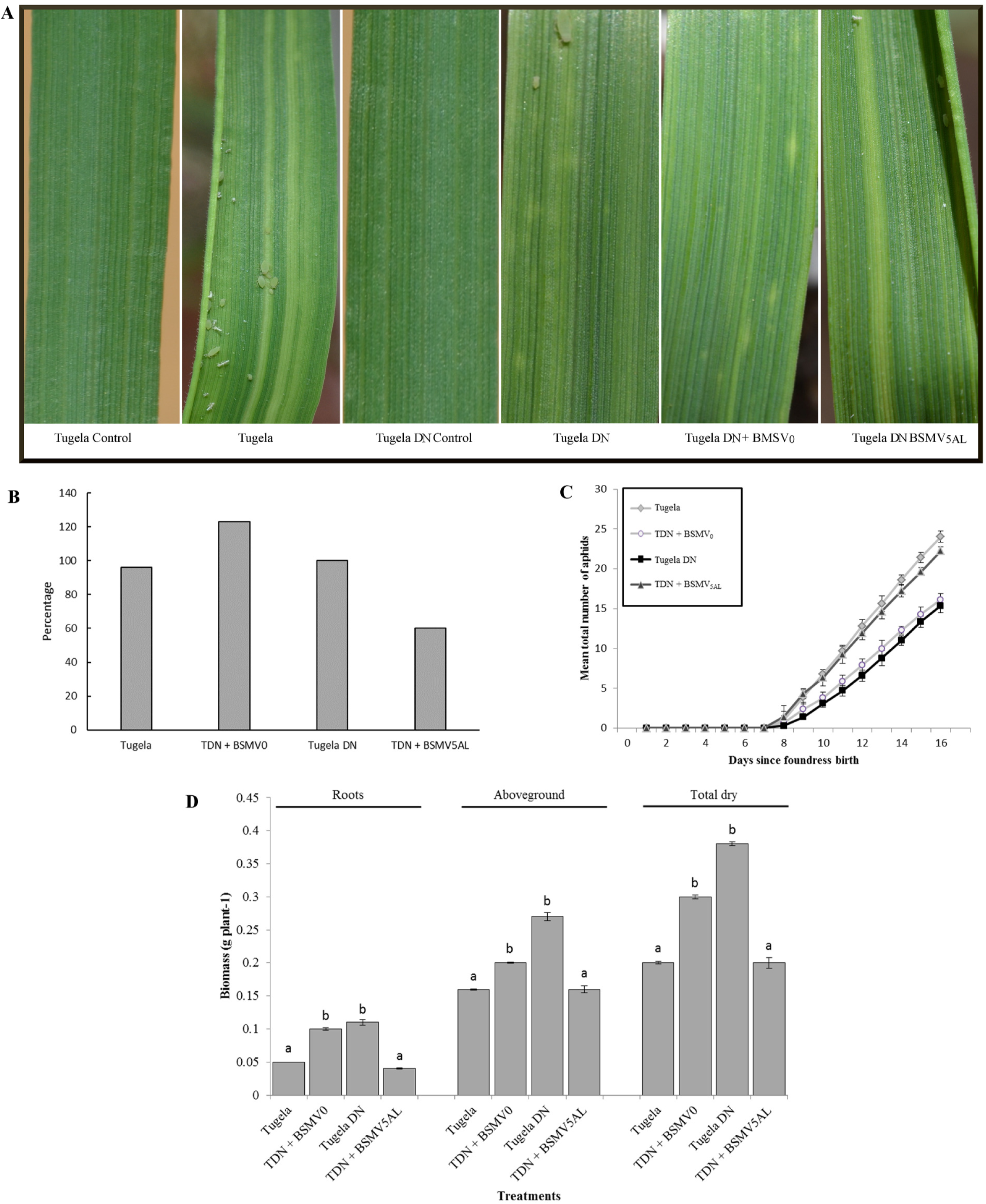
Plant and aphid phenotypic responses to BSMV-mediated VIGS of *5AL-B4.* **A**, Phenotypes of representative plants per treatment 19 days after viral infection. Both resistant controls (Tugela DN and TugelaDN+BSMV0) exhibited localized cell death lesions compared to the susceptible (Tugela) and Tugela DN+BSMV5AL treated plants that exhibited longitudinal chlorotic streaks characteristic of RWASA-1 infestation on susceptible cultivars. Tugela and Tugela DN Control represent uninfested plants, and Tugela and Tugela DN infested plants. **B,** Percentage knock-down of *5AL-B4* measured using ddPCR. **C,** Mean total aphid production of ten plants per treatment over 16 days. Aphids feeding on Tugela and Tugela DN had the highest and lowest mean production of nymphs respectively with the number of nymphs produced on silenced plants comparable to those on susceptible plants. **D,** Individual plant tissues and total dry plant biomass for each treatment following 16 days of aphid feeding on each treatment. Resistant controls had significantly higher biomass for all components compared to the susceptible control and the Tugela DN+BSMV_5AL_ treated plants.

Silencing of *5AL-B4* in Tugela DN induced phenotypic symptoms similar to the susceptible Tugela. The observed localized cell death lesions were replaced by yellowing and longitudinal chlorotic streaking similar to infested susceptible Tugela (Fig. 5A). This indicated that lower levels of *5AL-B4* resulted in chlorophyll loss similar to when *D. noxia* feeds on susceptible wheat plants. The above and belowground plant biomass was monitored to determine the effects of *5AL-B4* silencing on the health of the plant (Fig. 5E). The above-, belowground, and total dry weight of TDN+BSMV5AL-B4 was similar to the susceptible control Tugela, and significantly less (p = 0.0011) than that of the Tugela DN and TDN+BSMV0 controls. The severe reduction of plant biomass following a compatible infestation with *D. noxia* was consistent with previous reports (Anderson et al. 2008; Mirik et al. 2009). The reduced accumulation of plant biomass in silenced plants together with an increased *D. noxia* reproductive ability correlated to an inability by the plant to initiate an antibiotic defense response characteristic of *Dn1* mediated resistance.

## Discussion

Here we show that silencing of *5AL-B4* confers a susceptible phenotype to resistant Tugela DN upon *D. noxia* infestation. This is evident by increased aphid numbers, lowered plant vigor,loss of localized cell death and increased chlorotic streaking. The resistance mechanism in wheat plants carrying the *Dn1* resistance gene against *D. noxia* has been phenotypically characterized as antibiosis. This is typified by the host plant reducing aphid fecundity and adult aphid longevity (Botha et al. 2005). The TDN+BSMV*_5AL-B4_* plants did not show any alteration in the prenymphipositional period of the aphids, but an increase in aphid fecundity compared to resistant controls was evident. *Diuraphis noxia* infestation on susceptible hosts causes the breakdown of chloroplast and cellular membranes, leading to longitudinal chlorotic streaking (Botha et al. 2005). In resistant hosts, infestation leads to the development of small necrotic lesions similar to the localized cell death response observed in the HR (Botha et al. 2006; van Ooijen et al. 2008). TDN+BSMV_*5AL-B4*_ plants displayed inhibited formation of necrotic lesions around feeding sites and increased yellowing and extensive chlorotic streaking of the leaves. Tugela DN plants are known to produce reactive oxygen species (ROS), specifically H202, which activates downstream defense genes and is known to have a strong signaling and defense role during an incompatible interaction in plants with antibiotic resistance (Moloi and Van der Westhuizen 2006; Van Eck et al. 2010). The production of H_2_0_2_ is also necessary to induce salicylic acid accumulation via an increase in benzoic acid-2 hydroxylase activity (Léon et al. 1995), a defense mechanism against *D. noxia* in plants that harbor the *Dn1* resistance gene (Botha et al. 2005). In *Dn7* containing plants, which also confers an antibiotic effect on the plant, silencing of *phenylalanine ammonia-lyase (PAL)* and *WRKY53* disrupted the production of H202. This led Van Eck et al. (2010) to postulate that PAL and WRKY53 function as part of the defense cascade downstream of *Dn7.* Taken together, these data could indicate that *5AL-B4* functions close to, or possibly at, the molecular recognition of *D. noxia* by plants harboring the *Dn1* resistance gene.

WRKY integrated fusions are observed across plant lineages and are considered to represent recurrent fusions of the WRKY domain in diverse hosts. However, Sarris et al. (2016) suggests that an ancient fusion event of an AtWRKY46 homolog was integrated into monocotyledonous species prior to the divergence between wheat, sorghum, barley, and foxtail millet. The divergence between panicoid species (such as sorghum) and pooid species (such as wheat and barley) occurred approximately 40 million years ago (Murphy 2011), with the split from oryzoid species occurring earlier at 50 million years ago (Bossolini et al. 2007). Our phylogenetic analysis of all the known *T. aestivum* WRKY containing proteins indicated that the *5AL-B4* integrated WRKY domain is most closely related to TaWRKY50, whilst the C-terminal WRKY domain present in the homoeolog 5DL-AA14 is identical to TaWRKY41. A closer inspection of the integrated WRKY domains in other monocotyledonous species clearly indicated their unrelatedness to AtWRKY46, but that they rather cluster together within the panicoid, pooid, oryzoid, and dicot lineages (supplementary Fig. 1). This supports the recurrent integration of different WRKY domains for each lineage rather than a single ancient integration event prior to the divergence of the monocotyledonous species as previously proposed. The integration of different WRKY domains onto NLR genes could correlate with the NLR function with evolutionary pressure driving diversification of the WRKY domain within each host.

The WRKY integration into the *5DL-AA14* homoeolog is nearly identical to the WRKY integration in *Aegilops tauschii* and *Hordeum vulgare,* indicating a very well conserved NLR-ID following their divergence eight million years ago (Middleton et al. 2014). The *5AL-B4* NLR-ID is comparatively much younger, with its conserved ANK-CC-NLR sequence present in *T. urartu.* This precursor to *5AL-B4* was conferred to the hexaploid wheat progenitors during a hybridization event 0.2 – 1.3 million years ago when *T. urartu* hybridized with an unidentified B genome species to form tetraploid *T. dicoccoides* (Middleton et al. 2014). The WRKY domain was integrated once the complete hexaploid genome was formed approximately 8,000 – 10,000 years ago (Middleton et al. 2014) as it has no close relatives in any of the progenitor species. Within *T. aestivum* it is assumed to have evolved from a TaWRKY50 protein its closest relative. It still remains unclear why the *5AL-B4* integrated WRKY has diverged to such an extent, compared with the highly conserved WRKY domain in its homoeolog 5DL-AA14. We cannot rule out that the WRKY transcription factor *5AL-B4* recruited its domain from, could since have been lost from the wheat genome. However, this scenario is highly unlikely as there is no close homolog other than TaWRKY50 to the integrated WRKY on *5AL-B4* in any of the wheat progenitors. This would indicate that the WRKY would have had to be born and died within a very short span of time. Why *5AL-B4* has an evolved WRKY domain that does not closely match a WRKY transcription factor is under further investigation.

The role of TaWRKY50 and its nearest homolog in barley (HvWRKY21) is unclear, although some information is available on a close *Arabidopsis* homolog, AtWRKY41 (81.7 % homology). AtWRKY41 is a flagellin induced gene involved in the incompatible interaction between *Arabidopsis* and the biotrophic pathogen *Pseudomonas syringae* pv. *tomato.* Overexpression of AtWRKY41 increases resistance towards *Pseudomonas* but decreases the resistance towards the necrotrophic pathogen *Pectobacterium carotovorum* (Higashi et al, 2008). *Pseudomonas syringae* suppresses AtWRKY41 expression through a type III secretion system effector in compatible interactions (Higashi et al. 2008). Sarris et al. (2016) subsequently found an NLR-ID with an AtWRKY41 domain that interacts with the effector AvrRps4 from *Pseudomonas* in a yeast-two hybrid screen. Based on the integration of a similar domain onto *5AL-B4*, it is conceivable that these two proteins function in a similar manner, and we propose that *5AL-B4* could be functioning in effector trapping and are currently investigating this.

Sarris et al. (2016) found fourteen NLR-IDs occurring as double fusions, where a protein kinase domain was fused with an additional domain, either sequentially or each domain separated by the NLR core. The integration of more than one domain onto the NLR-ID may serve to detect more than a single effector or perhaps one of the two domains may have biochemical activity while the second domain simply detects the effector. These proteins appear to have developed from sequential fusion events, as is most probably the case with *5AL-B4.* The closest relative of *5AL-B4* is an ankyrin repeat domain containing NLR from the wheat “A” genome progenitor *T. urartu.* Unlike *5AL-B4,* this protein does not contain an integrated WRKY domain and is 94 % homologous to *5AL-B4* if the non-homologous C-terminals following the LRR are not considered. These two proteins are the only ankyrin repeat domain containing NLRs that have been identified from extensive database searches. This indicates that a unique fusion event occurred in *T. urartu* that integrated the ankyrin onto the NLR with subsequent donation to the *T. aestivum* genome. A second fusion event integrated the WRKY domain onto *5AL-B4* to create a unique NLR-ID protein within the plant kingdom. Interestingly, the P-loop in the ancestral form of *5AL-B4* in *T. urartu* is missing (Fig. 1B). The conserved p-loop motif in the NB-ARC domain regulates nucleotide binding and mutations within this motif abrogate the ability of the NLR to confer disease resistance or activate the HR (van Ooijen et al. 2008), which indicates that the ancestral gene may not have been functional and subsequent reactivation occurred in *5AL-B4.* Additionally, the functionality of the ancestral gene may have been lost after its donation to *Triticum.*

The amino-terminal domain of plant NLRs may be involved in both the detection of the pathogen signal and activation of downstream signaling molecules (DeYoung and Innes 2006; Collier and Moffett 2009). While the majority of NLR proteins contain a TIR or CC-domain at their N-termini, some proteins have no sequence N-terminal to the NB-ARC domain. A small number of proteins may have a Solanaceae domain or a BED DNA binding domain replacing or in conjunction with the CC-domain (Collier and Moffett 2009). Ankyrin repeat domain-containing proteins constitute one of the largest protein families in all species and plays a role in protein recognition and binding (Mosavi et al. 2004; Vo et al. 2015). Proteins contain one to 33 repeats, although at least two repeats are necessary in order to assume a folded structure (Leila et al. 2004). In plants, ankyrin domain-containing proteins are involved in a wide variety of biological processes with the majority linked to defense responses (Vo et al. 2015). Observations of ankyrin protein-protein interactions involved in plant defense show that they bind and perceive effectors in the plasma membrane, cytosolic signal transduction and activation of nuclear defense gene expression, depending on the subcellular localization signal of the ankyrin repeat domain-containing protein (Vo et al. 2015). In the case of *5AL-B4,* the ankyrin repeats may mediate intermolecular interactions, where they bind proteins involved in the defense gene cascade once the effector has been detected by the WRKY motif and the receptor has been activated. It can also not be ruled out that they are actively recognizing and interacting with effectors themselves.

Sequencing of *5AL-B4* from the resistant and the susceptible lines revealed no SNPs or indels between them or compared to the cv. Chinese spring that would account for a resistant genotype. Focusing on the increase in aphid fecundity and absence of H_2_0_2_ in TDN+BSMV_*5AL-B4*_ plants, *5AL-B4* plays a role in the defense response of wheat against *D. noxia,* suggesting an interaction with an unidentified binding partner to form a receptor complex much like previously reported NLR-ID genes. This binding partner is either absent or mutated in the susceptible line compared to the resistant line.

*5AL-B4* contains domains not found in its closest progenitors. This could be the result of unequal crossing over or gene conversion as seen in other NLR genes that occur as neighbors at the same locus (Loutre et al. 2011). Thus, these paralogs can serve as sources of variation. However, like *Lr10* and *RGA2, 5AL-B4* is a single copy gene that has no closely related NLR genes at its locus leaving it without access to variation generation due to crossing over with other paralogs. As no other close relative of the WRKY domain, with TaWRKY50 being the closest, was identified in the wheat genome it is not clear where the WRKY domain originated. It could be a result of gene conversion and allelic recombination between ancient haplotypes or that divergence of the close relatives preceded duplication (du Preez et al. 2008) and this would be in accordance with loss of genes after genome duplication events as observed in polyploids (Blanc and Wolf 2004). A counter argument to this would be that the short timespan might not have allowed for this to have occurred.

This is the first report of a WRKY containing NLR-ID protein that functions during a plant-pest interaction. The role of *5AL-B4* in this defense response is intriguing as it is not the *Dn1* gene, but could well be interacting with it directly. The *Dn1* gene was mapped to a different portion of the genome and Tugela and Chinese Spring do not have *D. noxia* resistance. Furthermore, there are no SNPs evident between the two alleles in Tugela and Tugela DN or between them and Chinese Spring. Thus, we postulate that *5AL-B4* is a NLR-ID protein that is needed to dimerize with *Dn1* to facilitate resistance in Tugela DN and propose the name *Associated with Dn resistance 1 (Adr1)* to reflect the loss of *Dn1* resistance once it has been silenced. Additionally, silencing of the WRKY-containing *5ALB4* and WRKY53 (Van Eck et al. 2010) might be indicative that *D. noxia* has developed analogous effectors to phytopathogens that target the evolutionary conserved WRKY transcription factors functioning in innate immunity.

## Materials and Methods

### Plant and insect growth conditions

All experiments were performed using two near isogenic wheat lines (NILs), Tugela (RWASA1 susceptible) and Tugela DN (PI 137739 – *Dn1,* RWASA1 resistant) that were obtained from the Agricultural Research Council – Small Grain Institute (ARC-SGI, Bethlehem, South Africa). The resistant Tugela DN was created by back-crossing the *Dn1-*gene from SA1684 into Tugela. This gene has been mapped to the 7DS chromosome in Tugela DN (Bierman 2015). The plants were grown to the two-leaf stage (Zadoks stage 12) under controlled conditions at 18 °C with a 12 h photoperiod for approximately fourteen days after germination for all experimental procedures. The RWASA1 aphids were obtained from the ARC-SGI and maintained on commercially available susceptible PAN3434 wheat plants (Pannar Seeds, Greytown, South Africa) under controlled conditions at 18 °C with 12 h photoperiods before use. The three repetitions of barley stripe mosaic virus (BSMV) inoculations were performed on 14 plants per treatment, with uninoculated Tugela and Tugela DN used as the susceptible and resistant controls respectively.

### Sequence and phylogenetic analyses

TRIAE_CS42_5AL_TGACv1_374266_AA1195550 (November 2014 *T. aestivum* Ensemble release; designated *5AL-B4* from here) is an NLR with similar architecture to NLR-ID decoys against pathogen effectors. It shared homology with *Pi36* that was identified as a target for differentially regulated miRNA from the study of Nicolis et al. (2017) and identified as a possible role player in the wheat-*D. noxia* interaction. This, coupled with its unique architecture, prompted us to further its role in the wheat-*D. noxia* interaction. The full length of the gene was amplified from both NILs using cDNA and cloned into pGEM-T Easy (Promega) for sequence analysis. The amplification was performed using 250 nM of each of three primer sets (Set1-F: 5’-CCGGAAATGTTGCCCTTGTG-3’; Set1-R: 5’-CATAGCACGGTCTTCCGCTCTC-3’; Set2-F: 5’-GCCACGTCCACATGCTTCCTAG-3’; Set2-R: 5’-GACGAACCTTGTCTGCGAGTG-3’; Set3-F: 5’-TCCTGCACACTGCATCACATGG-3’; Set3-R: 5’-ACGCGCTGACATCAAATTCG-3’) that spanned the length of the gene using KAPA HiFi HotStart (KAPA Biosystems). The sequences for both NILs were generated and aligned to TRIAE_CS42_5AL_TGACv1_374266_AA1195550 to identify polymorphisms. Sequences downloaded from Ensemble were aligned using MAFFT and a phylogenetic analysis was performed using maximum parsimony analysis with PAUP* version 4b10 (Swofford et al. 2002). A maximum likelihood analysis was performed using PhyML to ascertain the placing of *5AL-B4* using standard parameters and set to determine the best fit model. Bootstrap support (1,000 replicates) were determined for the tree’s branching points. The consistency and retention indices were determined for all the datasets. Full-length sequences were used to determine the placing of *5AL-B4* and only the WRKY domains were considered to identify the closest WRKY relative of *5AL-B4*.

### Expression analyses

Confirmation of the differential regulation for *5AL-B4* in the wheat-*D. noxia* interaction was established using droplet digital (ddPCR). The expression for *5AL-B4* was studied at nine time points (0, 0.5, 1, 2, 4, 6, 8, 24, 48 hpi) in both NILs. Each plant was infested with 20 aphids, allowed free movement, and non-infested controls were included as reference points for gene expression. RNA was extracted from five wheat plants per treatment for three biological repeats using the Plant RNeasy Mini extraction kit (Qiagen). The RNA from the five extractions per sample were mixed in equimolar concentrations and a total of 1 µg RNA was converted into cDNA using iScript (Bio-Rad) and used at a 1:19 dilution as template in a ddPCR reaction containing 2X QX200 ddPCR EvaGreen Supermix (Bio-Rad) and 200 nM of primer B4-F (5’-TCCTGCACACTGCATCACATGG-3’) and B4-R (5’-GACGAACCTTGTCTGCGAGTG-3’). Reaction droplets were generated in a QX200 Droplet generator (Bio-Rad) using a DG8 cartridge and the PCR was performed on a T100 thermal cycler (Bio-Rad) using a ramp rate of 2 °C s^-1^ and enzyme activation step at 95 °C for 5 min. This was followed by 40 cycles of 95 °C for 30 s and 60 °C for 1 min. A final signal stabilization step of 4 °C for 5 min followed by 90 °C for 5 min was performed. Data acquisition was performed on a QX200 droplet reader and data analyzed using QuantaSoft Software (Bio-Rad).

### Virus induced gene silencing of *5AL-B4*

The sequence following the WRKY domain in *5AL-B4* is unique. An across-species and wheat specific BLASTn revealed no potential silencing of non-target transcripts. From this unique sequence a 270 bp fragment was amplified from cDNA using the VIGS-F (5’- ACACGTGCTTGGACTCTGTC-3’) and VIGS-R (5’-CGAATTTGATGTCAGCGCGT-3’) primers. The fragment was amplified using KAPA HiFi HotStart ReadyMix, cloned into the pSL038-1 vector and verified through sequencing. The construction of the BSMV silencing vector and method of viral inoculation followed the protocol by Scofield et al. (2005). Viral controls included BSMV_0_, which is derived from the empty pSL038-1 vector, and BSMV_PDS_ that included a transcript that targets the *phytoene desaturase* gene and acts as a visual marker of correct viral reconstitution. Both Tugela and Tugela DN plants were treated with the virus constructs to ascertain the effect of silencing *5AL-B4* on both NILs. Five days after viral inoculation, the plants were mass infested with 10 *D. noxia* apterous adults. Additionally, a single apterous aphid was placed inside a clip cage attached to the emergent third leaf to determine the fecundity of the aphids feeding on the different silenced plants. The following day, all aphids apart from one new born nymph were removed from the cage. The remaining new born was regarded as the foundress and nymphs born to this foundress were counted and removed every 24 h for 14 days. The intrinsic rate of increase (rm) for each foundress was estimated according to the equation r_m_ = (0.738 × In(M_d_))/d, where M_d_ is the number of nymphs produced in a period equal to the prereproductive time (d) (Wyatt and White 1971). To determine the integrity of the antibiosis defense mechanism in *Dn1* carrying Tugela DN, the effects of aphid feeding on plant biomass accumulation was assessed. Six days after mass aphid infestation (eleven days after viral inoculation), aphids were removed from the third leaf of three experimental plants per treatment and the leaf tissue collected into liquid nitrogen and stored at −80 °C prior to extraction. RNA was extracted from each individual leaf sample by homogenization in liquid nitrogen followed by purification with the RNeasy plant mini kit. Droplet digital PCR was used for absolute quantification of *5AL-B4* transcript levels in VIGS treated plants as described previously. Twenty one days after viral inoculation, all aphids were removed from three plants per treatment and the aboveground plant biomass was separated from the roots. The roots were rinsed and together with the aboveground plant biomass dried for 48 h at 40 °C and their weight determined.

## Acknowledgments

The authors would like to thank A Jacobs for assistance with the phylogenetic analyses and critical discussion on the manuscript; and, Bio-Rad for the use of the QX200 machine to quantify transcript abundance. This work was funded by public grants from the Winter Cereal Trust (WCT/W/2007/04 Induced systemic resistance in wheat) and the National Research Foundation Technology and Human Resources for Industry Programme (TP2009072000010 and TP2011070700029).

## Author contributions

VFN and EV contributed equally to the research design, data analyses, and preparation of the manuscript. Research funding was obtained by EV.

---

**Supplementary Fig. S1.**

Phylogenetic analysis of a representative number of WRKY domain containing NLR-IDs from monocotyledonous species against AtWRKY46. Analysis indicates that multiple, lineage and host specific WRKY integration events occurred.

